# A whole-organ multi-scale in silico framework for human kidney haemodynamics informed by hierarchical phase-contrast tomography

**DOI:** 10.64898/2025.12.30.696985

**Authors:** Yousef Javanmardi, Mengzhe Lyu, Yang Zhou, Yufei Wang, Claire L Walsh, Paul Tafforeau, Daniyal J. Jafree, David A. Long, Alexandre Bellier, Ryo Torii, Peter D. Lee

**Affiliations:** Department of Mechanical Engineering, University College London, UK; Siemens Healthineers Digital Technology (Shanghai) Co. Ltd., Shanghai 200124, China; European Synchrotron Radiation Facility, Grenoble, France; Developmental Biology and Cancer Research and Teaching Department, UCL Great Ormond Street Institute of Child Health, London, UK; UCL Centre for Kidney and Bladder Health, University College London, London, UK; Wellcome Trust Sanger Institute, Hinxton, Cambridge, UK; University of Grenoble Alpes, Department of Anatomy (LADAF), CIC INSERM 1406, Grenoble, France; Research Complex at Harwell, Rutherford Appleton Laboratory, Didcot, UK

## Abstract

Studying human kidney haemodynamics has been limited by the absence of complete vascular maps of the whole organ. Here we utilise previously generated multi-resolution hierarchical phase-contrast tomography (HiP-CT) coupled with a hybrid anatomically-grounded synthetic reconstruction to generate a full arterial–glomerular network of an intact human kidney comprising 1.6 million vessels and over 800,000 glomeruli. Using this anatomically comprehensive structure, we apply physics-based zero-dimensional haemodynamic modelling to quantify blood pressure, flow and simulated filtration rate across the entire organ. We show that the reconstructed human kidney vascular network exhibits order-dependent branching behaviour similar to that of rat kidneys, and that physiologically plausible pressure and flow patterns are recovered only when the full vascular network is represented. We further demonstrate how the kidney responds to macrovascular and microvascular perturbations, including stenosis of the large renal arteries and narrowing or ablation of afferent arterioles. Stenosis and arteriole narrowing exhibit threshold-type behaviour, with kidney perfusion and simulated glomerular filtration rate remaining largely preserved up to ∼50% narrowing, followed by sharp nonlinear declines beyond ∼70%. These predictions emerge in the absence of autoregulatory mechanisms, indicating that vascular geometry and resistance scaling alone contribute to kidney functional deterioration. Together, our framework provides the first organ-wide, data-driven model of human kidney haemodynamics and offers a foundation for future studies of kidney physiology and disease.

## Introduction

Kidney function relies on a sophisticated, multiscale vascular network^1^ that delivers blood from the renal artery to nearly one million nephrons^2^. The kidney arterial tree forms a hierarchical branching structure in which parent vessels progressively divide into multiple generations of daughter vessels, producing a characteristic pressure drop from proximal to distal branches^3^. At the terminal level, afferent arterioles supply the glomerular capillary tufts where plasma is filtered, a process estimated clinically as the glomerular filtration rate (GFR), a surrogate indicator of overall kidney function^4^. Understanding how vascular branching architecture shapes pressure distribution, flow division, and ultimately glomerular filtration is central to our understanding of kidney physiology in health and disease.

Although blood pressure and flow can be measured *in vivo*, such measurements are accessible only at limited locations and cannot characterise haemodynamics across the entire kidney vascular network^5^. Assessment of multiscale kidney haemodynamics can therefore be achieved using *in silico* modelling, which provides a controlled framework for analysing pressure and flow across the vascular hierarchy^6^. These quantities are accessible *in vivo* only at limited locations and not measurable across the full kidney vascular network. Three-dimensional computational fluid dynamics (3D-CFD) resolves detailed flow patterns in individual vessels but requires prohibitively large computational resources at the whole organ scale^7^. In contrast, zero-dimensional (0D) lumped-parameter models represent each vessel as an equivalent resistor^8^, enabling efficient simulation of whole-kidney pressure and flow distributions^9^ and facilitating model-based estimation of GFR^10^. This 0D approach has been applied in rodents, offering insights into how branching structure influences flow heterogeneity across vessel generations^11–13^. However, haemodynamic modelling of the human kidney has been limited primarily by the absence of anatomically complete vascular data^6,14^. High-resolution techniques such as micro-computed tomography (CT)^15^ and confocal imaging^16^ can visualise microvasculature but only within small tissue volumes^17^, whereas clinical CT^18^, MRI^19^, and angiographic modalities^20^ capture only a few generations of the entire vascular network.

Recent advances in hierarchical phase-contrast tomography (HiP-CT)^21–23^, a synchrotron X-ray imaging technique, provide a unique multiresolution imaging capability, enabling whole-organ visualisation while preserving micron-scale detail in selected regions. In our previous work, we used HiP-CT to obtain an image-derived arterial tree containing nearly 10,000 vessels across nine Strahler generations^14^, a substantial improvement over conventional imaging and the most complete human kidney vascular map available to date. However, even with this level of detail, the smallest vascular branches remain below the whole-organ imaging resolution. To address this limitation, several strategies have been developed, including rule-based methods that use experimentally derived statistical laws^24,25^ and image-based approaches that segment the resolvable arteries directly from CT or micro-CT data^26^. Although each approach has its advantages, a hybrid strategy offers a more comprehensive solution^11,27^: this approach preserves the anatomically accurate large vessels obtained from imaging while generating the unresolved small vessels as statistically inferred branches, thereby completing the pre-glomerular network in a subject-specific manner.

Here, we utilise a hybrid strategy to generate a complete vascular network of a structurally intact human kidney which we then integrate with a 0D lump-parameter model to simulate whole-kidney haemodynamics. Arteries from the largest down 8-9 generations are taken directly from our prior HiP-CT data^14^, while the smallest vascular branches are added as statistically inferred segments. Each afferent arteriole is coupled to a physiologically grounded terminal resistance derived from dedicated 3D CFD simulations of glomerular haemodynamics, allowing us to obtain model-based estimates of simulated GFR (sGFR). After establishing this integrated model, we first examine haemodynamics under physiological conditions. Next, we assess the impact of limited imaging resolution by systematically truncating the vascular network to mimic the reduced number of vessel generations accessible in clinical imaging. Finally, we evaluate pathological scenarios including stenosis of the large renal arteries as well as afferent-arteriole narrowing or loss. Together, these analyses provide mechanistic insight into the structural determinants of kidney haemodynamics and function, with relevance to health and disease.

## Results

### Hybrid multiscale reconstruction of the human renal arterial network

To reconstruct the full kidney arterial network, we first applied an image-based approach using HiP-CT to capture the larger (> 30 μm), well-resolved vessels of a structurally intact human kidney (**Figure 1a**). The HiP-CT dataset was obtained from a previously analysed sample^14^ (LADAF-2021-17), a 62-year-old male donor with no documented kidney disease (cause of death: pancreatic cancer). The organ was imaged at 25 µm voxel size and binned to 50 µm for whole-organ analysis^28^. As shown in **Figure 1b**, this resolution enables clear delineation of the major arteries and their branching patterns throughout the entire organ. In our previous study^14^, we segmented, skeletonised, and quantified this arterial tree, yielding 10,189 image-derived arterial segments spanning nine Strahler generations (**Figure 1c, SI Figure 1**). These image-derived vessels form the structural backbone of the hybrid reconstruction, capturing the anatomically accurate geometry of the large and medium-sized arteries.

**Figure 1.**
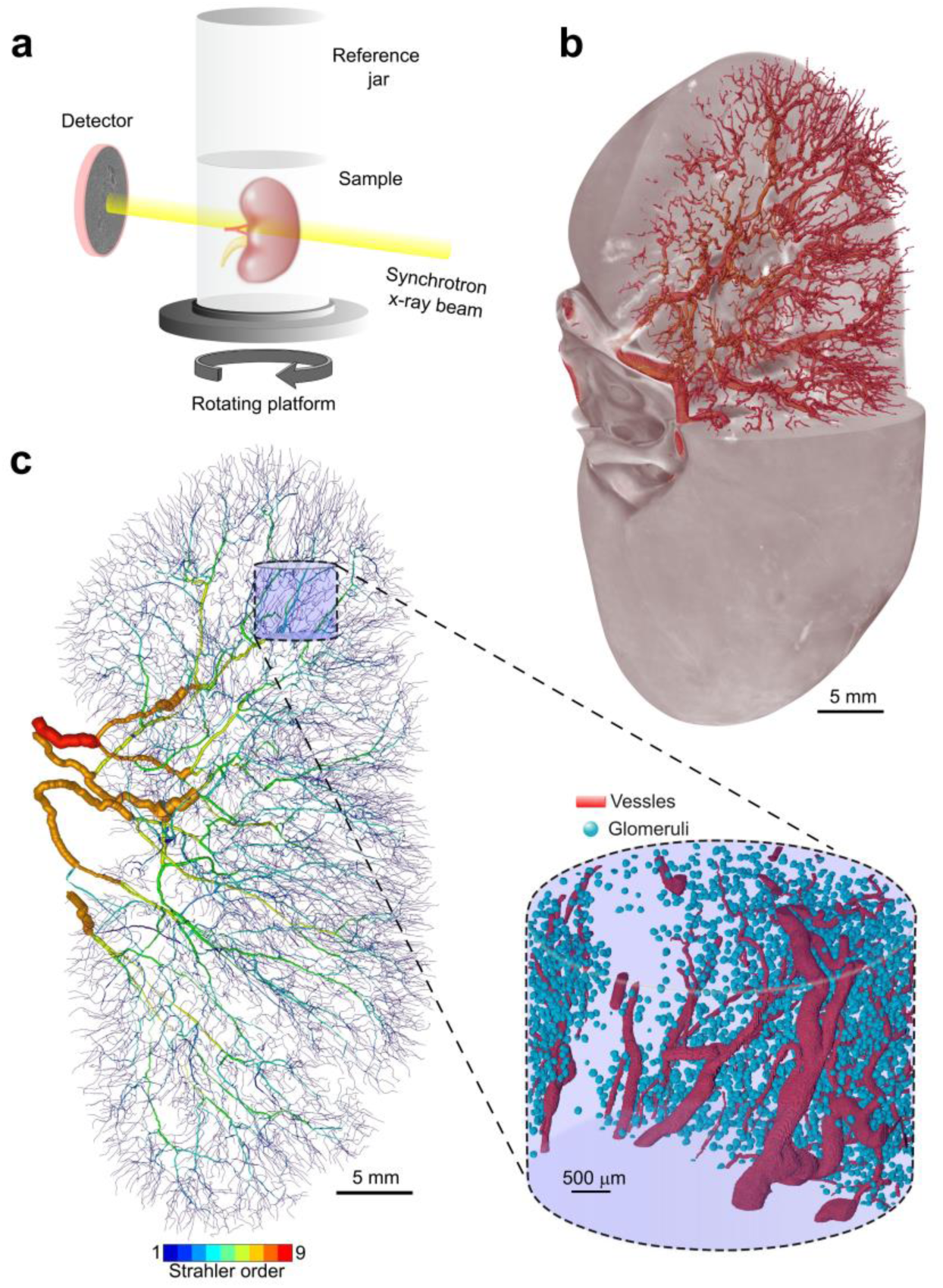
Multiresolution HiP-CT imaging of the human kidney and extraction of image-derived arterial and glomerular structures. **(a)** Schematic of the Hierarchical Phase-Contrast Tomography (HiP-CT) acquisition setup, in which the intact kidney is rotated in a synchrotron X-ray beam while projections are recorded on a detector. **(b)** Whole-organ rendering of the kidney imaged at 25 µm voxel size (resampled to 50 µm for analysis), overlaid with the segmented arterial network obtained in our previous work^14^. **(c)** Skeletonised arterial tree from the 50 µm dataset, colour-coded by Strahler order (1–9), with vessel thickness proportional to diameter. The highlighted mauve region marks the location of a high-resolution HiP-CT zoom scan acquired at 2.6 µm and binned to 5.2 µm. The inset shows segmented arteries and glomeruli within this zoom region, used for deriving statistical laws for unresolved vessels and for mapping glomerular endpoints.

To characterise the small vessels unresolved at 50 µm resolution, we analysed a high-resolution HiP-CT zoom region of the same kidney, imaged at 2.6 µm voxel size^29^ and binned to 5.2 µm for segmentation (**Figure 1c**). The process of inferring statistical rules for synthesising the missing branches consisted of two stages. First, to derive empirical laws for expanding the 50 µm arterial tree down to the smallest resolvable vessels, we segmented and skeletonised the arterial network in the 5.2 µm zoom dataset (**Figure 2a**), identifying 294 vessels within this sub-volume. When the same region was examined in the 50 µm whole-organ dataset, only 52 vessels were present (**Figure 2a**), revealing the scale of under-representation of distal branches at whole-organ resolution.

**Figure 2.**
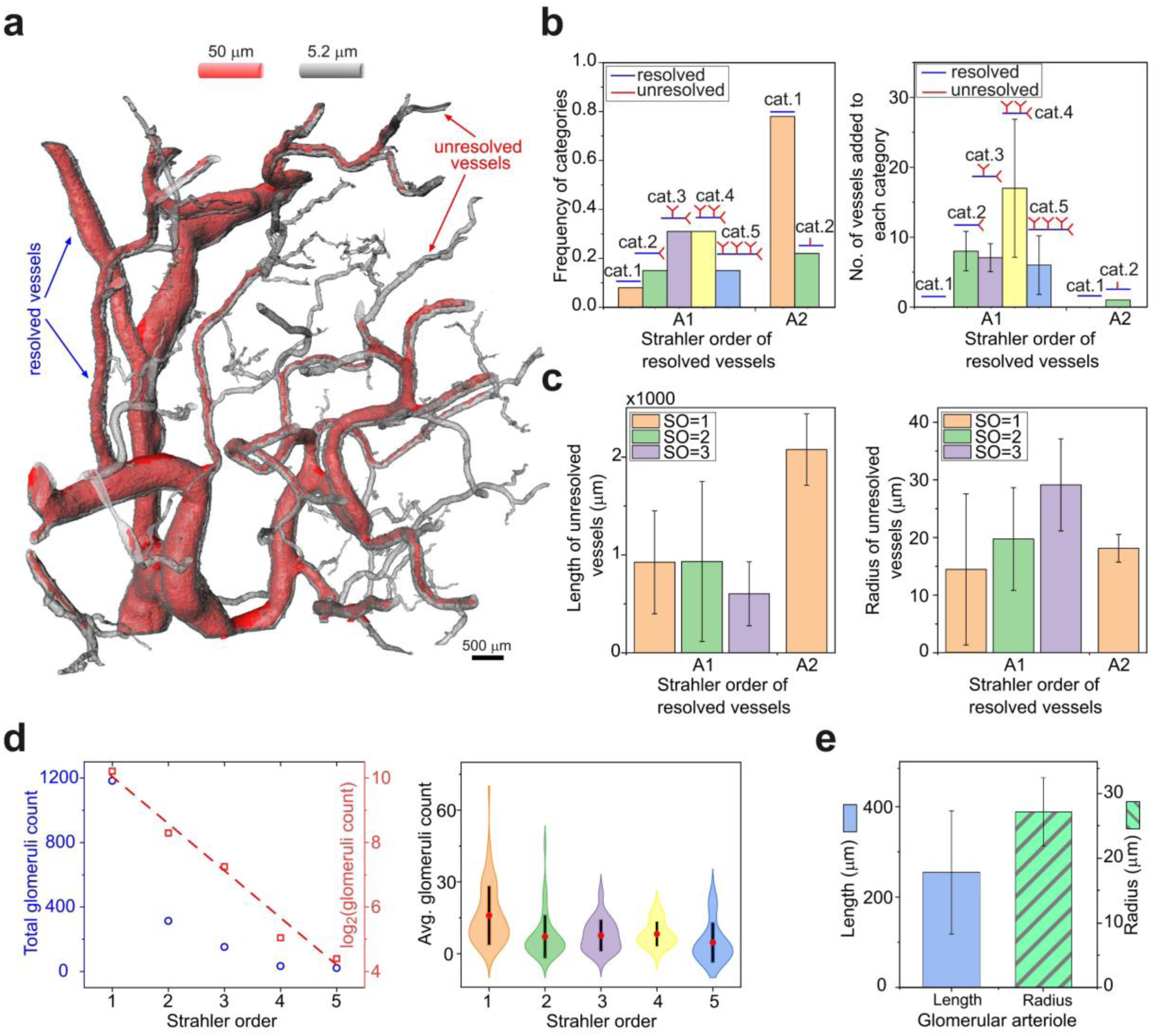
Statistical inference of unresolved pre-glomerular vessels and glomerular assignment based on high-resolution HiP-CT data. **(a)** Comparison of the arterial network resolved at 50 µm and 5.2 µm voxel size in the zoom region. Vessels visible at both resolutions are labelled as *resolved vessels* (blue), while vessels visible only at 5.2 µm but absent in the 50 µm dataset are labelled as *unresolved vessels* (red). **(b)** Classification of unresolved vessels arising from each resolved parent vessel. Left: frequency of unresolved-vessel categories for Strahler-order 1 (A1) and Strahler-order 2 (A2) parent vessels. Right: total number of unresolved vessel segments added per category. Schematic icons illustrate each category. **(c)** Length and radius of unresolved vessel segments stratified jointly by (i) the Strahler order of their *resolved parent vessel* in the 50 µm dataset (A1 or A2) and (ii) the Strahler order assigned to the unresolved segments themselves in the 5.2 µm reference tree (SO = 1, 2, or 3). **(d)** Distribution of glomeruli extracted from the high-resolution reference dataset. Left: total glomerular count per Strahler order shown on linear and log₂ scales. Right: kernel density estimates of the number of glomeruli associated with each Strahler order. **(e)** Length and radius distributions of afferent arterioles, extracted from those resolvable at 2.6 µm resolution and used to generate synthetic afferent arterioles throughout the kidney. All error bars show mean ± s.d.

Direct comparison of the two datasets showed that the additional vessels found in the 5.2 µm dataset predominantly arise from arteries classified as Strahler order 1 (A1), with a smaller contribution from Strahler order 2 (A2), in the 50 µm tree. We therefore quantified, for each A1 and A2 vessel, the probability and branching pattern of the additional subtrees required to match the high-resolution anatomy found in the 5.2 µm zoomed scan. This analysis revealed that approximately 8% of A1 vessels required no additional inferred segments (Category 1). About 15% required the addition of two terminal subtrees, comprising a total of eight vessel segments (Category 2). A further 31% required the addition of two terminal subtrees and one side subtree, together comprising seven vessel segments (Category 3). Following the same observational approach, three additional categories (Categories 4-6) were defined as shown in **Figure 2b**. The geometric characteristics of these synthetic subtrees, including the length and radius of their constituent vessel segments, are shown in **Figure 2c**. Applying these empirically derived rules across all A1 and A2 vessels in the 50 µm network resulted in the addition of 800,091 new arterial segments, producing a multiscale network that captured substantially more of the vascular architecture of the kidney. However, this expansion still did not resolve the afferent arterioles leading to glomeruli, which remain too thin and tortuous to be visualised even at 5.2 µm resolution.

Therefore, we incorporated glomeruli and their corresponding afferent arterioles to complete the distal portion of the arterial network. For this purpose, we used a set of glomerular positions obtained previously through a deep learning–based segmentation pipeline developed by Zhou et al.^30^, which was applied to the same 5.2 µm HiP-CT zoom region. By mapping each glomerulus to its nearest arterial segment in the high-resolution reference tree, we quantified the distribution of glomeruli per Strahler order (**Figure 2d**) and used this to assign glomeruli across the entire kidney, resulting in a total of 813,926 glomeruli. Because afferent arterioles could be visualised only in a small number of cases at 2.6 µm due to their tortuosity and small calibre, we extracted their length and radius distributions from the identifiable examples (**Figure 2e**) and applied these distributions generatively across all glomerular endpoints. Together, these steps yielded a complete vascular network with arterial–glomerular connectivity, comprising 1,624,206 vessel segments distributed across 12 Strahler generations, forming the anatomical basis for whole-organ haemodynamic modelling.

### Organ-scale 0D simulation of human kidney haemodynamics

Having reconstructed the anatomically complete vascular network spanning from the main renal artery to glomerular endpoints (**Figure 3a**), we next simulated whole-organ kidney haemodynamics using a 0D lumped-parameter model. Here, Strahler order 12 corresponds to the main renal artery, which serves as the model inlet where pressure or flow boundary conditions are imposed. At the opposite end of the hierarchy, Strahler-1 segments represent the afferent arterioles, each of which terminates at a glomerulus (**Figure 3a**); in total, 813,926 glomeruli were incorporated as terminal points.

**Figure 3.**
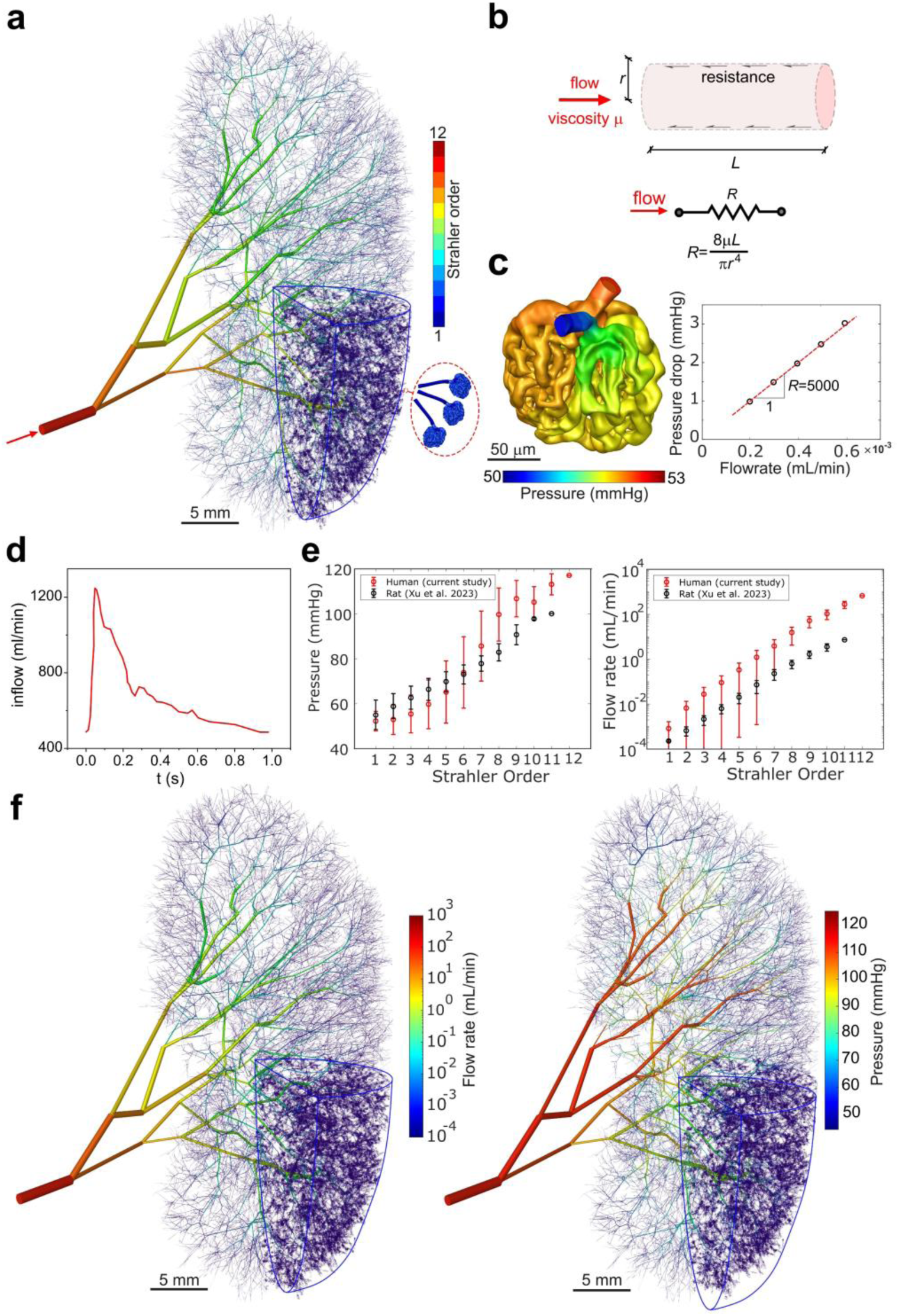
Organ-scale 0D simulation of renal haemodynamics in the reconstructed human kidney. **(a)** Fully reconstructed multiscale arterial network containing 1,624,206 vessel segments across 12 Strahler orders, with Strahler-1 terminals corresponding to afferent arterioles connected to 813,926 glomeruli. **(b)** Schematic of the 0D lumped-parameter modelling framework, in which each vessel segment is represented by a hydraulic resistance determined by its length, radius, and blood viscosity. **(c)** Determination of glomerular resistance using 3D CFD simulations performed on a detailed human glomerulus geometry provided by Terasaki et al.^31^, yielding a linear pressure–flow relationship. **(d)** Pulsatile flow profile measured in vivo by Doppler ultrasound in the human renal artery (Krejza et al.^32^), used as the model inlet boundary condition. **(e)** Simulated pressures and flow rates as a function of Strahler order in the reconstructed human kidney, compared with the order-dependent trends reported for rat kidneys by Xu et al.^11^ (error bars show s.d.). **(f)** Spatial distribution of mean pressure and flow throughout the reconstructed arterial tree. Strahler-1 vessels are omitted outside the blue-outlined region for clarity and improved visibility of global flow patterns.

In this model, each vessel segment is represented by a hydraulic resistance determined by its length, radius, and blood viscosity (**Figure 3b**). Under the assumption of laminar flow, the pressure drop across a segment is linearly related to the flow passing through it, as described by Poiseuille’s law. Mass conservation at each bifurcation ensures that the sum of daughter flows matches the parent flow, allowing pressures and flow rates to be determined throughout the entire network once inlet and terminal boundary conditions are specified. This formulation provides a computationally efficient means of propagating pressure and flow through more than 1.6 million reconstructed vessels.

Within the 0D framework, the glomerulus is treated as an effective hydraulic resistance, analogous to individual vessel segments. Because the glomerular capillary meshwork lacks an analytical pressure–flow description, we used a detailed 3D reconstruction of a representative human glomerulus provided by Terasaki et al.^31^ and performed CFD simulations to quantify its pressure drop over physiological flow rates (**Figure 3c**). The resulting linear pressure–flow relationship was used to define an effective glomerular resistance, which was then added in series to the resistance of each corresponding afferent arteriole to capture both pre-glomerular and intraglomerular pressure losses.

Boundary conditions were set using an *in vivo* renal artery flow waveform measured by Doppler ultrasound in humans by Krejza et al.^32^, in which flow varies between 440 and 1270 mL/min over one cardiac cycle thus a cycle-average of 667 mL/min was derived (**Figure 3d**). Using this inflow condition, the model predicted a progressive pressure drop from ∼115 mmHg at the main renal artery (Strahler order 12) to ∼57 mmHg at Strahler order 1 (**Figure 3e**), following an approximately linear (Pearson *r* = 0.979, R^2^ = 0.96) trend across orders. This pattern is consistent with the order-dependent pressure decline reported in rat kidneys reported by Xu et al.^11^ Flow rates showed a similar order dependence: on a semi-logarithmic scale, flow decreased linearly (Pearson *r* = 0.995, R^2^=0.99) with decreasing Strahler order (**Figure 3e**). The spatial distribution of flow rate and pressure across the whole kidney is shown in **Figure 3f**. Notably, the flow map (**Figure 3f**, left) closely mirrors the Strahler-order anatomy shown in **Figure 3a**, reflecting the approximately linear relationship between the logarithm of flow rate and Strahler order. Together, these results demonstrate that human kidney haemodynamics exhibit order-dependent patterns resembling those reported previously in rat kidneys.

### Effect of incomplete vascular reconstruction on whole-kidney haemodynamics

Clinical imaging resolves only the largest renal arteries, typically capturing just a few Strahler orders^33,34^, and haemodynamic models based on such data therefore rely on substantially incomplete vascular trees, as in Qi et al.^35^. To assess how this limited resolution affects haemodynamic predictions, we systematically truncated the reconstructed human arterial tree by removing successive distal generations and repeated the 0D simulations on each pruned network (**Figure 4a**). This approach quantifies how excluding small vascular branches, those commonly unresolved in clinical scans, alters predicted pressure and flow distributions.

**Figure 4.**
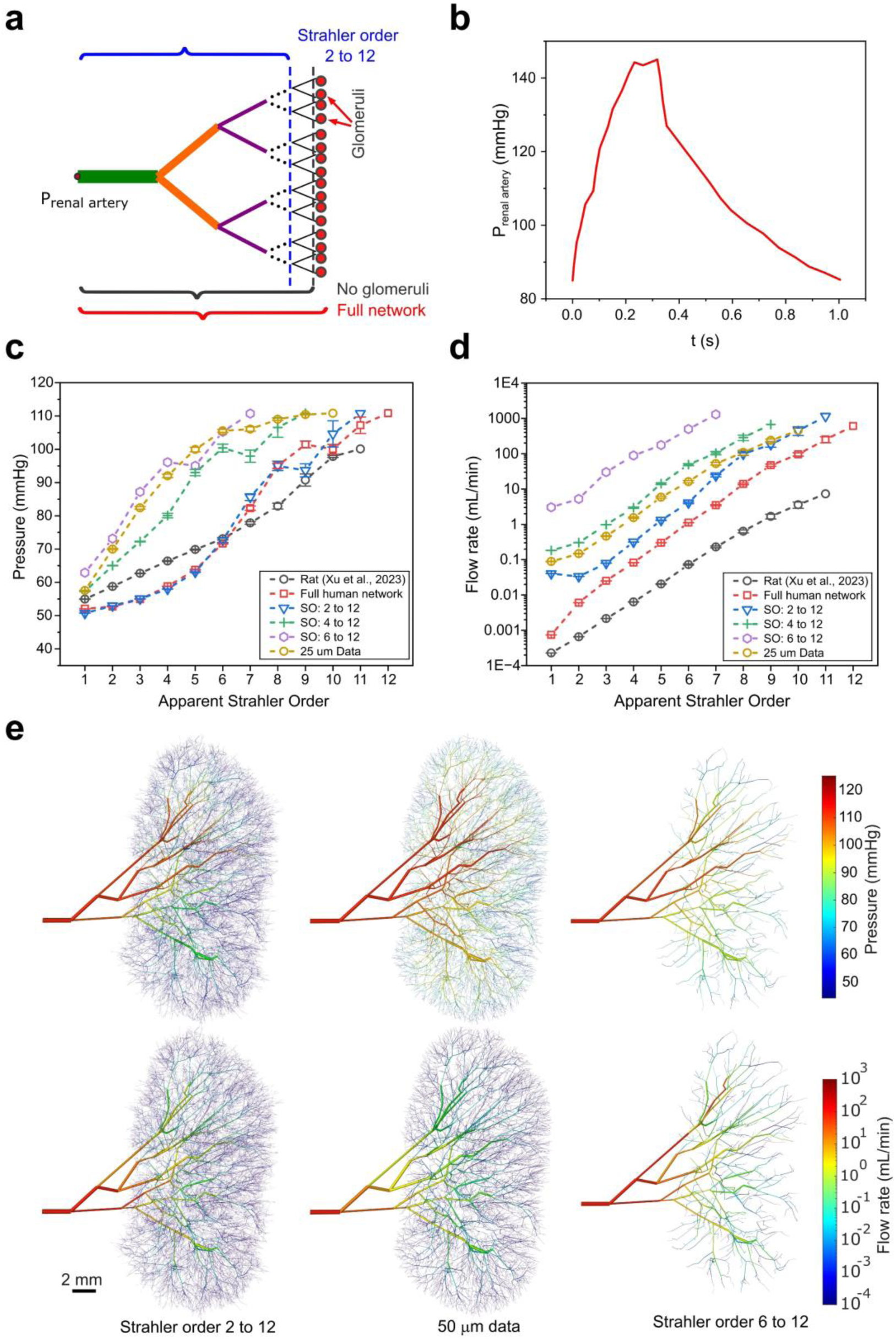
Impact of vascular tree truncation on renal pressure and flow. **(a)** Schematic illustration of the pruning approach, in which distal vessel generations were systematically removed to mimic the limited resolution of clinical imaging. **(b)** Pulsatile inlet pressure profile applied at the renal artery, taken from Carter et al.^36^. **(c)** Simulated pressure as a function of Strahler order for the full network and for networks truncated at different distal orders, compared with rat data from Xu et al.^11^. **(d)** Corresponding flow rates for the same pruning conditions. (Error bars show standard error) **(e)** Spatial maps of pressure (top row) and flow (bottom row) for three representative cases: pruning Strahler order 1 (left), using all vessels resolved at 50 µm (middle), and pruning Strahler orders 1–5 (right).

A pulsatile pressure boundary condition measured *in vivo* by Carter et al.^36^, ranging from ∼85 to 145 mmHg (mean ∼107 mmHg), was applied at the renal artery inlet (**Figure 4b**). Progressive truncation of distal vessel generations led to a systematic increase in pressure within mid-level branches (**Figure 4c**). For example, at Strahler order 6, the average pressure increased from ∼73 mmHg in the full network to ∼71, 93, 97, and 105 mmHg when vessels of orders 1–5 were sequentially removed (**Figure 4c and SI Figure 2a, c**). This behaviour arises because the total pressure drop across the kidney remains fixed by the pressure boundary conditions, and removing distal generations shortens the number of levels over which this drop must occur. Consequently, the pressure difference assigned to each remaining generation increases, steepening the pressure–Strahler order slope.

Flow rates also increased with successive pruning (**Figure 4d**). At Strahler order 6, mean flow rose from ∼1.1 mL/min in the complete network to 1.7, 4.0, 17.4, 49.0, 110.4, and 498.7 mL/min as orders 1–5 were removed (**SI Figure 2b, d**). This occurs because, with the pressure boundary conditions fixed, reduced vascular resistance simply increases flow. Spatial patterns of pressure and flow for representative pruning levels are illustrated in **Figure 4e**, where the loss of distal generations leads to progressively higher pressures and larger flows in the remaining branches. These results also suggest a practical application for clinical imaging scenarios in which only a limited number of vascular generations are resolved and the pressure boundary conditions at terminal endpoints are unknown. In such cases, pressures predicted by the full-network simulations at these endpoints could be used as physiologically informed boundary conditions when modelling flow in truncated vascular networks derived from clinical images.

Together, these findings demonstrate that incomplete vascular reconstruction leads to systematic, order-dependent distortions in predicted kidney pressure and flow, with errors increasing as distal generations are progressively excluded.

### Applying *in silico* perturbations to the kidney vasculature and modelling their effects on organ-wide haemodynamics

#### Proximal inflow restriction

Having established organ-scale haemodynamics in the intact human kidney, we next used the model to examine how structural perturbations alter renal perfusion. We focused first on proximal inflow restriction, introduced by imposing graded narrowing (0–90%) of the radius at the main renal artery (**Figure 5a**). This setup mimics conditions in which inflow restriction leads to reduced downstream perfusion, such as renal artery stenosis, embolus, or surgical clamping^37^. The inlet pressure boundary condition, derived from *in vivo* measurements^38^, ranged between ∼86 and ∼162 mmHg and was based on guidewire recordings from patients (n=18, age range 27-85 years, 61% male) with renal artery stenosis (**SI Figure S3a**).

**Figure 5.**
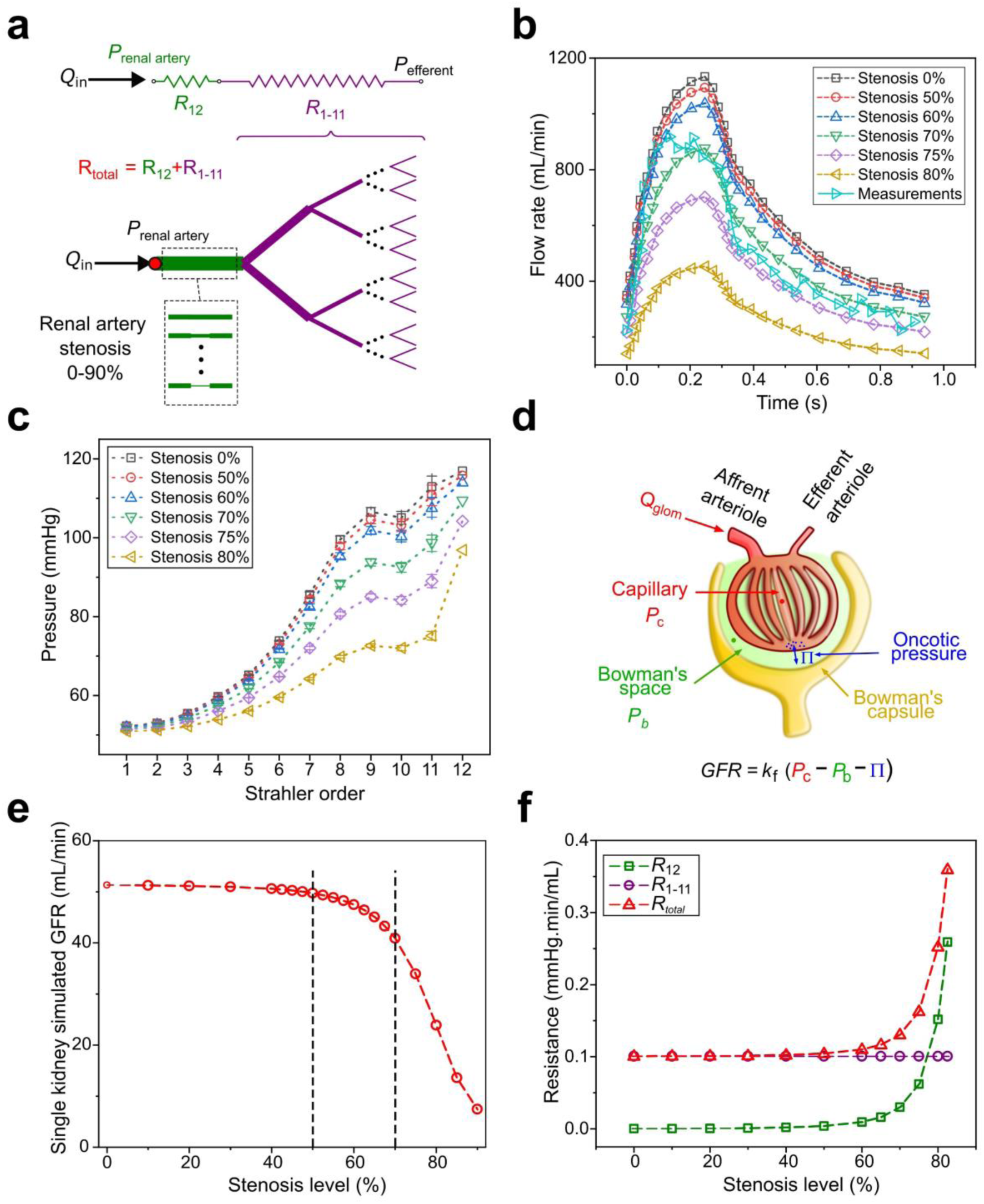
Modelling renal artery stenosis as a cause of hypoperfusion and its impact on haemodynamics and filtration. **(a)** Schematic illustrating the implementation of renal artery stenosis in the model (0–90% narrowing) and the associated decomposition of total renal resistance into the resistance of the stenotic segment (R₁₂) and the downstream vascular network (R₁–₁₁). **(b)** Simulated renal artery flow waveforms for different stenosis levels compared with in vivo measurements from guidewire recordings (taken from Collard et al.^38^). **(c)** Mean intrarenal pressure as a function of Strahler order under increasing degrees of stenosis. **(d)** Illustration of the filtration model used to compute single-nephron simulated GFR (sGFR), in which glomerular capillary pressure (P_c_) from the 0D simulation drives filtration opposed by Bowman’s space pressure (P_b_) and oncotic pressure (π). **(e)** Simulated whole-kidney glomerular filtration rate (sGFR) across stenosis severities, showing preserved filtration up to moderate narrowing and a sharp decline beyond ∼70–80% stenosis. **(f)** Changes in segmental resistances with stenosis severity, showing the increase in R₁₂ relative to the relatively stable downstream resistance R₁-₁₁, and the resulting increase in total renal vascular resistance (R_total).

Simulations showed increasing stenosis severity progressively reduced both peak and mean renal blood flow at the inlet (**Figure 5b**). Peak flow in the model without stenosis reached ∼1130 mL/min, whereas among the simulated conditions, the 70% stenosis case most closely aligned with the clinically reported flow waveform^38^ (**Figure 5b**), suggesting that, despite studying a different kidney, our model captures characteristic haemodynamic responses to proximal inflow restriction. Strikingly, pressure distributions across the vascular tree remained remarkably stable at stenosis levels up to 50%, with mean pressures at all Strahler orders deviating only modestly (< 2%) from baseline (**Figure 5c**). Similarly, total renal blood flow also declined only slightly over this range (from ∼665 mL/min at baseline to ∼642 mL/min at 50% stenosis). At higher severities, however, both pressure and flow decreased substantially. For example, the pressure at Strahler order 11 fell from 112 mmHg in the intact model to 73 mmHg at 80% stenosis, and total renal blood flow dropped to ∼266 mL/min (**SI Fig. S3b**).

To estimate consequences on filtration, we used the intrarenal pressures predicted by the 0D model to compute the pressure within each glomerular capillary (Pc), which provides the driving force for filtration. We then applied a filtration model based on Fick’s principle (**Figure 5d**) and calculated the single-nephron filtration rate as a function of the pressure drop across each glomerulus. Summing across all nephrons in a kidney yielded a model-derived estimate of single-kidney filtration, referred to here as the sGFR. The model predicted that sGFR remained stable at ∼51 mL/min up to 50% stenosis. A mild reduction (∼7%) appeared at 70% stenosis, followed by a sharp decline at higher severities, with an ∼85% reduction at 90% stenosis (**Figure 5e**). These findings highlight the haemodynamic resilience of the kidney to moderate reductions in inflow pressure and suggest that glomerular function is largely preserved until hypoperfusion exceeds a critical threshold.

Stenosis caused a progressive increase in the resistance of the proximal renal artery (R₁₂): rising from 0.0002 mmHg·min/mL in the healthy case to 0.003 mmHg·min/mL at 50% stenosis, and reaching ∼0.03 mmHg·min/mL by 70% stenosis. In contrast, the resistance of the downstream network (R₁–₁₁) remained constant at ∼0.1 mmHg·min/mL. As a result, total vascular resistance increased only modestly for stenosis levels below 70%, but rose sharply once R₁₂ exceeded R₁–₁₁, reaching ∼0.35 mmHg·min/mL at 85% stenosis and producing the observed nonlinear escalation in total resistance (**Figure 5f**).

#### Distal microvascular stenosis or ablation

Having evaluated how proximal inflow changes affect kidney haemodynamics, we next investigated how perturbations to the distal microvasculature, specifically the afferent arterioles, alter pressure and flow distribution across the kidney. We implemented two approaches, (i) removing a proportion of afferent arterioles (and their associated glomeruli), simulating loss of functioning nephrons as observed in chronic kidney disease and ageing^39–41^ (**Figure 6a-d**), and (ii) reducing afferent arteriolar diameter, mimicking functional narrowing observed in primary hypertension^42^, or certain nephrotoxins such as non-steroid anti-inflammatory drugs^43,44^ (**Figure 6e-h**). These complementary perturbations allowed us to examine how nephron-level microvascular loss redistributes pressure and flow throughout the arterial tree and to quantify its impact on kidney function by computing the sGFR.

**Figure 6.**
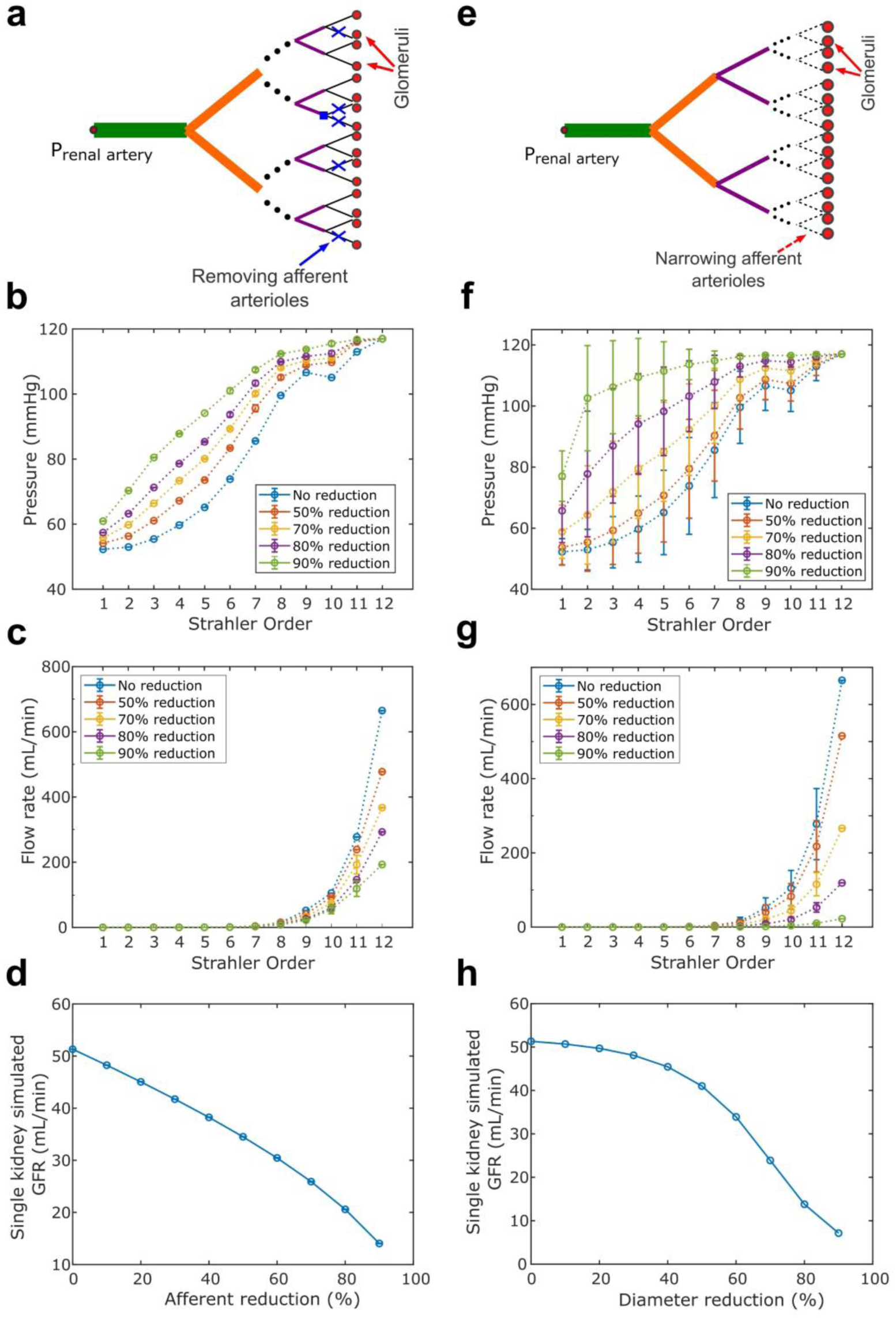
Haemodynamic and functional consequences of distal microvascular perturbations. **(a)** Schematic showing afferent arteriole ablation: random removal of 10–90% of afferent arterioles and their glomeruli in 10% increments (*n* = 10 independent simulations per level). Nodes with no remaining downstream afferents were closed (blue squares). **(b)** Arterial pressure across Strahler orders for each removal level. **(c)** Flow rate across Strahler orders for each removal level. **(d)** Simulated single-kidney GFR (sGFR) as a function of the percentage of afferent arterioles removed. **(e)** Schematic showing afferent arteriole narrowing: reducing afferent arteriole diameter by 10–90% in 10% increments. **(f)** Arterial pressure across Strahler orders for each diameter-reduction level. **(g)** Flow rate across Strahler orders for each diameter-reduction level. **(h)** sGFR as a function of the percentage reduction in afferent arteriole diameter.

To model afferent arteriole ablation, we applied the pressure boundary conditions indicated in **Figure 4b** and progressively removed 10–90% of afferent arterioles in 10% increments. For each selected arteriole, both the vessel and its corresponding glomerulus were removed (**Figure 6a**). Unlike the analysis in Figure 4, when all afferent arterioles downstream of a branching node were removed, the node was closed, preventing any further outflow through that subtree (blue square in **Figure 6a**). This progressive truncation produced a marked rise in mid-level arterial pressures. For example, mean pressure at Strahler order 6 increased from 74 mmHg in the intact network to 83, 89, 94, and 101 mmHg when 50%, 70%, 80%, and 90% of afferent arterioles were removed, respectively (**Figure 6b**). Correspondingly, total renal inflow decreased substantially, with flow at the main renal artery falling from 665 mL/min (no reduction) to 477, 367, 293, and 193 mL/min across the same ablation levels (**Figure 6c**). Interestingly, both pressure elevation and flow decline were intensified beyond ∼50% ablation. The resulting functional impact was captured by sGFR, which decreased approximately linearly from 51 to 26 mL/min between 0–70% ablation, followed by a steeper decline to 14 mL/min at 90% removal (**Figure 6d**).

To model afferent arteriole narrowing (stenosis), vessel diameters were reduced by 10–90% in 10% increments while applying the same pressure boundary conditions as in **Figure 4b**. Progressive narrowing led to substantial increases in mid-level arterial pressure: at Strahler order 6, mean pressure rose from 74 mmHg in the intact kidney to 79, 92, 103, and 114 mmHg at 50%, 70%, 80%, and 90% diameter reduction, respectively (**Figure 6f**). Concomitantly, total renal inflow declined sharply, with flow at the main renal artery decreasing from 665 mL/min to 515, 266, 119, and 23 mL/min across the same degrees of narrowing (**Figure 6g**). Strikingly, the impact on kidney function exhibited a pronounced bi-linear pattern: sGFR decreased modestly from 51 to 41 mL/min up to 50% narrowing, after which further constriction caused a steep decline, reaching 7 mL/min at 90% reduction (**Figure 6h**).

Together, these results show that kidney haemodynamics and filtration are resilient to low-to-moderate proximal inflow restriction and distal microvascular perturbations, whereas more severe perturbations at either the main renal artery or the level of afferent arterioles are associated with pronounced pressure redistribution, reduced inflow, and a marked decline in sGFR.

## Discussion

Understanding human kidney haemodynamics has long been constrained by the absence of complete, organ-scale vascular maps. Here we leveraged a hybrid anatomical reconstruction based on multiresolution HiP-CT to obtain a full multiscale arterial network of an intact human kidney. This network provided the foundation for physics-based 0D haemodynamic modelling, allowing us to simulate pressure, flow, and filtration rate across the entire organ. Together, this framework reveals how vascular geometry shapes intrarenal perfusion and enables mechanistic exploration of clinically relevant perturbations, such as those observed in different forms of kidney disease, ageing or when the kidney is exposed to certain drugs that modify haemodynamics.

Here, we analysed a kidney from a 63-year-old donor with no documented renal disease and reconstructed the full arterial–glomerular network using our hybrid HiP-CT–based approach. This yielded ∼814,000 glomeruli, consistent with biopsy-based estimates from large donor cohorts of similar age^45^. For the haemodynamic simulations, we applied a renal artery flow waveform measured in healthy individuals^32^ (**Figure 3d**), which produced a main renal artery pressure of ∼115 mmHg—well within the normal clinical range^46^. We then compared the resulting human data with a detailed rat dataset and found that both species share the same hierarchical branching pattern. In both species, the number of vessels decreases exponentially with Strahler order, appearing as a straight line on a log₂ scale (**SI Figure S2a**). Because flow across each branching point must be conserved, this exponential branching causes the input flow to be progressively divided among an increasing number of daughter vessels towards distal vessels, resulting in the linear relationship between log flow and Strahler order observed in both datasets (**Figure 3e**). A similar order dependence is seen for pressure, which increases systematically from distal to proximal orders in both humans and rats (**Figure 3e**). Notably, these relationships hold only when the vascular tree is complete; using the 50 µm image-derived network alone produces an inaccurate, bilinear pressure–order profile (**Figure 4c**). This emphasises the need for high-resolution, multiresolution imaging such as HiP-CT to capture the full pre-glomerular architecture and obtain reliable haemodynamic predictions.

Next, we examined how limited inflow, by constriction of main renal artery, affects renal haemodynamics and the resulting sGFR (**Figure 5**). In the absence of stenosis, the model predicted an sGFR of ∼51 mL/min, consistent with *in vivo* observation for a healthy human single kidney in individuals of similar age^45,47,48^. As stenosis increased, sGFR exhibited a clear bilinear dependence on diameter reduction: it remained largely stable up to ∼50% narrowing, declined modestly at moderate stenosis (50–70%), and then dropped sharply when stenosis exceeded ∼70% (**Figure 5e**). This behaviour aligns with clinical classifications of renal artery stenosis severity—mild (<50%), moderate (50–70%), and severe (>70%)^49^—and mirrors clinical measurements, where single-kidney GFR typically decreases from 45±13 mL/min in mild to ∼34±17 mL/min in moderate stenosis and 19±11 mL/min in severe disease^50^. Notably, this threshold response emerged despite the absence of autoregulatory mechanisms in our model, indicating that arterial geometry and resistance scaling alone could explain the characteristic nonlinear decline in filtration. This can be understood by considering the relative contributions of upstream (R_12_) and downstream (R_1-11_) resistance (**Figure 5f**): at low to moderate stenosis, resistance of the main renal artery remains small compared with that of the remainder of the network, but once lumen reduction becomes sufficiently severe, renal artery resistance exceeds downstream resistance, causing a rapid rise in total vascular resistance and an abrupt fall in flow rate and glomerular pressure—ultimately producing the sharp decline in sGFR.

Finally, we examined how distal microvascular perturbations, modelled either as ablation of afferent arterioles and their associated glomeruli or as narrowing of the afferent arteriole lumen, alter whole organ haemodynamics. Both perturbations reflect clinically relevant pathophysiology: loss of afferent arterioles and nephrons accompanies ageing^51^ and progression of chronic kidney disease^52^, whereas afferent arteriole narrowing is commonly observed in hypertension^53^, diabetic nephropathy^54^, and drug-induced renal vasoconstriction^55^. However, the organ-level haemodynamic consequences of these perturbations remain poorly understood, largely because human afferent arteriolar networks cannot be resolved in vivo using existing imaging modalities. Clinical measurements indicate that reductions in GFR during nephron loss often follow a biphasic pattern^56^, with GFR largely preserved up to approximately 50% nephron loss^57^ and a rapid decline at more advance stages^57^. Experimental and clinical studies of afferent vasoconstrictive states similarly suggest a nonlinear, threshold-like reduction in GFR^58–60^. Our model captures this behaviour in the case of afferent arteriole narrowing, where lumen reduction beyond ∼50% produced a sharp fall in sGFR, whereas afferent arteriole loss did not exhibit a comparable threshold-like response, possibly reflecting the absence of physiological regulatory mechanisms such as autoregulatory control or vessel wall compliance.

Together, these results provide a mechanistic picture of human renal haemodynamics across multiple perturbations. However, several methodological limitations should be acknowledged. First, the use of an *ex vivo* fixed kidney may not fully reflect in vivo vascular geometry, as tissue fixation can lead to partial lumen collapse and altered vessel compliance; inflation or perfusion-based preparation before imaging may help preserve physiological morphology in future studies. Second, although the flow model incorporates non-Newtonian viscosity, it remains a single-phase, rigid-wall approximation and therefore cannot capture the full complexity of blood rheology or vessel mechanics. The reconstruction procedure also relies on statistical rules derived from a single high-resolution subregion. Moreover, our analysis is based on a single human kidney, and broader inter-individual variability cannot be assessed. Finally, the model does not incorporate autoregulatory mechanisms, which limits its ability to reproduce compensatory responses observed in vivo, particularly during structural nephron loss.

This study demonstrates how combining multiresolution HiP-CT with physics-based haemodynamic modelling can generate a complete, organ-scale view of renal blood flow and filtration. By linking vascular geometry to perfusion patterns and perturbation responses, our framework provides a mechanistic foundation for investigating human renal haemodynamics under both physiological and pathological conditions. Continued development, including integration of autoregulation, additional donor data, and improved imaging fidelity, will further enhance its ability to support clinical interpretation and the study of kidney disease.

## Materials and Methods

### Sample Preparation, HiP-CT Acquisition and Reconstruction

The sample preparation, HiP-CT acquisition, and reconstruction procedures are fully described in Rahmani et al.^14^. Briefly, a human kidney from a 63-year-old male donor (LADAF-2021-17)^28,29^ with no documented renal disease was formalin-fixed, embedded in agar, and imaged at the European Synchrotron Radiation Facility using Hierarchical Phase-Contrast Tomography (HiP-CT). Whole-organ imaging was performed at 25 µm voxel size (reconstructed at 50 µm after binning), and a cortical sub-region was acquired at 2.6 µm voxel size (reconstructed at 5.2 µm). Tomographic reconstruction, volume stitching, and preprocessing were carried out following the standard HiP-CT pipeline described in Walsh et al.^21^ and Brunet et al^61^.

### Image Segmentation and Skeletonisation

Arterial segmentation of the 50 µm whole-kidney dataset was carried out using the semi-automatic Magic Wand tool in Amira/Avizo, as described in Rahmani et al.^14^, with manual refinement to isolate arterial regions. The same procedure was applied to the 5.2 µm magnified dataset to extract small pre-glomerular vessels. Centreline skeletons for both datasets were then extracted using the automated skeletonisation method of Rahmani et al.^14^, yielding node–segment graphs suitable for geometric analysis. These skeletons were subsequently cleaned to remove spurious branches and merged to correct fragmented segments. The final processed skeletons were exported as spatial graphs for vascular reconstruction and haemodynamic modelling.

### Hybrid Vascular and Glomerular Reconstruction

A complete pre-glomerular arterial network was generated by combining image-derived vessels from the 50 µm whole-organ dataset with statistically inferred small vessels derived from the 5.2 µm magnified region. Image-resolved vessels were retained as ground-truth geometry for the larger arterial generations, while radius, length, and branching statistics measured in the high-resolution subvolume were used to generate synthetic vessels for unresolved distal orders. Synthetic segments were inserted at terminal nodes of the 50 µm skeleton and recursively expanded until the full pre-glomerular hierarchy was reconstructed. In the proximal renal artery region, several large branches were only partially captured in the image-derived data; the most proximal vessels including main renal artery were removed during the donor organ preparation, and the missing vessels were synthetically generated in our model by extending the available segments to their expected junction point, and the diameter of the parent vessel was estimated using Murray’s law.

Glomerular segmentation was obtained from our previous work^30^, which provides centroid positions for all glomeruli in the magnified region. To associate each glomerulus with its supplying afferent arteriole, we mapped glomerular centroids to the nearest segment of the reconstructed arterial tree using a 3D Euclidean distance criterion. Each glomerulus was then assigned a unique afferent arteriole, and terminal vessels were extended, where necessary, to ensure a one-to-one correspondence between glomeruli and afferent segments. The resulting combined vascular–glomerular graph served as the anatomical basis for haemodynamic simulations.

### Flow modelling

Flow in the arterial tree was simulated using a 0D lumped-parameter model (LPM) based on our previous work^62^. Here, each segment in the vascular tree is represented by a resistor, representing the segment’s haemodynamic resistance, as shown in **Figure 3b**. This method assumes that the blood flow in each arterial segment is a fully developed axisymmetric incompressible laminar flow of a homogeneous fluid, and the pressure loss due to branching is negligible. Resistance (*R_k_*) of each vessel segment is defined, following a commonly-adopted Poiseuille’s law, as follows^63^:

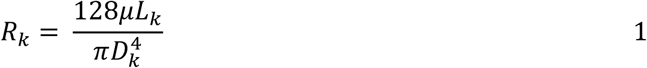

Here 𝐿_𝑘_ and 𝐷_𝑘_ are segment length and diameter of 𝑘-th vessel segment, respectively, 𝜇 is blood viscosity which depend on vessel diameter^24^:

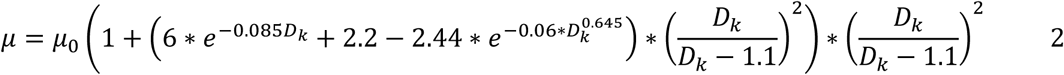

where 𝐷_𝑘_ here is the vessel diameter in micrometres and 𝜇_0_ = 1 𝑚𝑃𝑎. 𝑠 is a constant^64^. Given the large size of the dataset, direct simulation of flow throughout the entire renal vasculature with ∼1.6 million vessels was computationally infeasible due to memory constraints. To address this, we implemented a recursive calculation approach, in which calculation of flow and pressure is conducted in one Y-shaped bifurcation unit at a time.

The process starts with calculation of the equivalent resistance for entire renal vascular tree (i.e. 𝑅_12_ + 𝑅_1−11_ in **Figure 5**, note that 𝑅_1−11_ here includes glomerular resistances) by repeatedly adding the parallel and series resistances from the peripheral ends, based on the given vessel lengths and diameters. With the pressure at the main renal artery inlet and the end of vascular network (= 50 mmHg^65^), the flow into the entire vascular tree can be calculated (𝑄_1−12_). This can be split into the flows to the two sub networks of Strahler order 1-12 vessels 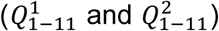 accordingly to the resistance of the respective sub network (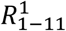 and 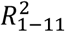), from which the pressure at the beginning of the sub network can be derived via 𝑃 = 𝑄𝑅 . Recursive execution of this process allows calculation of flow through each vessel segment and pressure at each vascular node throughout the network, i.e. 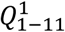 can be further split into 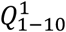 and 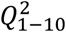, and 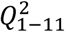 into 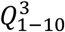 and 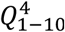, according to resistances of sub networks 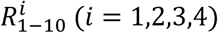.

This approach significantly reduced computational demands while maintaining an accurate representation of the extremely complex renal vascular anatomy, making it feasible to analyse large-scale datasets without exceeding memory limitations.

### Glomerular resistance estimation

To determine glomerular resistance, we utilised a three-dimensional (3D) glomerular structure obtained by Terasaki et al.^31^ through scanning electron microscopy (SEM). The dataset was provided as a surface model in .obj file format, which was subsequently processed for 3D computational fluid dynamics (CFD) simulations in ANSYS Fluent. The glomerular geometry was meshed to ensure accurate flow analysis, with a fine mesh resolution (average 0.4 μm) and inflation layers near boundaries to capture flow behaviour near the walls. To simulate physiologically relevant conditions, a downstream pressure of 50 mmHg^65^ was applied at the efferent arteriole, while the afferent arteriole pressure was varied in the range of 51–53 mmHg. This range was selected based on the known pressure drop across the glomerulus, which is typically only a few mmHg^47^. For each input pressure, the resulting flow rate was computed. The glomerular resistance was then determined as the slope of the pressure drop vs. flow rate curve (5000 mmHg/(mL/min)), which turned out to be a clear linear relationship in this physiological pressure range (**Figure 3c**). Rather than modelling glomeruli explicitly in the 0D flow simulation, the computed resistance value was incorporated into the resistance of all terminal vessels in the reconstructed vascular network.

A typical renal flow or pressure waveform obtained from the literature was applied at the inlet as shown in **Figures 3d** and **4b**. All outlet nodes, after efferent arterioles, where assigned a pressure of 50 mmHg as the boundary condition^66^.

### GFR estimation

The filtration model, depicted in **Figure 5d**, is based on models presented by Keener and Sneyd^67^. The filtration rate of a single glomerulus is estimated using the following equation^67^:

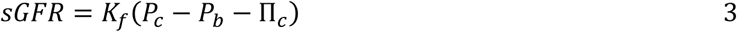

here, 𝐾_𝑓_ represents the glomerular ultrafiltration coefficient, Π_𝑐_ denotes the oncotic pressure in the plasma, and 𝑃_𝑐_ and 𝑃_𝑏_ are the average hydrostatic pressures in the capillary and Bowman’s capsule, respectively. This equation is based on the assumption that the rate of fluid loss from the capillary is linearly proportional to the pressure gradient as the driving force^68^. Capillary pressure (*P_c_*) was assumed to be the average pressure at the afferent and efferent arterioles. The afferent arteriole pressure right before the glomerulus was calculated using:

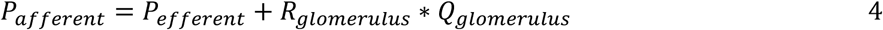

where *Q_glomerulus_* is the flow rate through the glomerulus, obtained from the 0D simulation. The pressure in Bowman’s capsule and oncotic pressure were assumed to be 15 mmHg^66^ and 34.75 mmHg^69^, respectively, while an efferent arteriole pressure of 50 mmHg was chosen^66^. A range of 5–50 nl/min/mmHg has been proposed for *K_f_* ^70^,; here, we selected an average value of *K_f_* = 27.5 nl/min/mmHg.

## ACKNOWLEDGEMENTS

This project has been made possible in part by the CZI grant 2022-316777 (DOI https://doi.org/10.37921/331542rbsqvn) from the Chan Zuckerberg Initiative DAF, an advised fund of Silicon Valley Community Foundation and the Royal Academy of Engineering (CiET1819/10). P.D.L. and C.L.W. gratefully acknowledge funding from the MRC (MR/R025673/1). P.D.L. is a CIFAR MacMillan Fellow in the Multiscale Human Program. D.J.J. is supported by a Wellcome Trust Accelerator Award (314710/Z/24/Z) and the Specialised Foundation Programme at the East of England NHS Deanery. D.A.L. is supposed by a Wellcome Trust Investigator Award (220895/Z/20/Z) and the NIHR Biomedical Research Centre at Great Ormond Street Hospital for Children NHS Foundation Trust and University College London. We gratefully acknowledge the European Synchrotron (ESRF) funding for beamtimes (md1252 & md1290), and all those who helped with them. The authors express their gratitude to Dr Mark Terasaki for sharing the 3D structure of a glomerulus.

