## Supplementary Information for "A whole-organ multi-scale in silico framework for human kidney haemodynamics informed by hierarchical phase-contrast tomography"

### Supplementary materials

Table S1. Metadata for image collection

| Donor | Pixel Size | Beamline | Energy (kEv) | Propagation Distance (mm) | Filters | Scintillator | No. of projections | Exposure Time (s) | Sensor name | Total scan time (hrs) |
|---|---|---|---|---|---|---|---|---|---|---|
| LADAF-2021-17 | 25.00 | BM05 | 81 | 3500 | Mo 0.1 SiO2 8x4mm | LuAG:Ce 1000um | 6000 | 0.035 | PCO edge 4.2 CLHS | 3.7 |

The body was embalmed by injecting 4500 mL of 1.15% formalin in lanolin followed by 1.44% formalin into the right carotid artery, before storage at 3.6 °C. During evisceration of the kidney, vessels were exposed, and surrounding fat and connective tissue removed. The kidney was post-fixed in 4% neutral-buffered formaldehyde at room temperature for one week. The kidney was then dehydrated through an ethanol gradient over 9 days to a final equilibrium of 70%.Each solution was four-fold greater than the volume of the organ and during dehydration, the solution was degassed using a diaphragm vacuum pump (Vacuubrand, MV2, 1.9m$^3$/h) to remove excess dissolved gas. The dehydrated kidney was transferred to a polyethylene terephthalate jar where it was physically stabilised using a crushed agar-agar ethanol mixture, and then imaged.

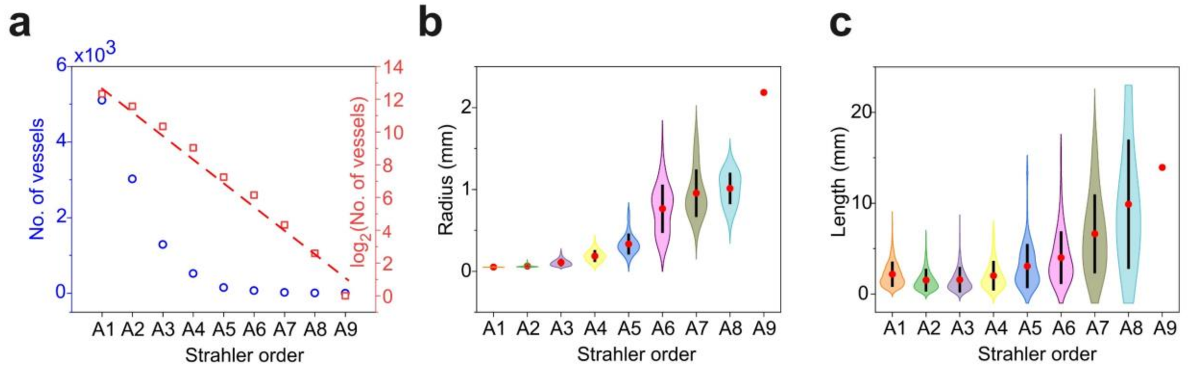


**SI Figure S1. Characterization of vessel properties obtained from 50 µm voxel size images (a)** Number of vessels per Strahler order, shown on linear and $log_2$ scales. **(b)** Vessel radius distributions across Strahler orders (mean shown in red). **(c)** Vessel length distributions across Strahler orders (mean shown in red).

### Impact of Incomplete network on pressure and flowrate

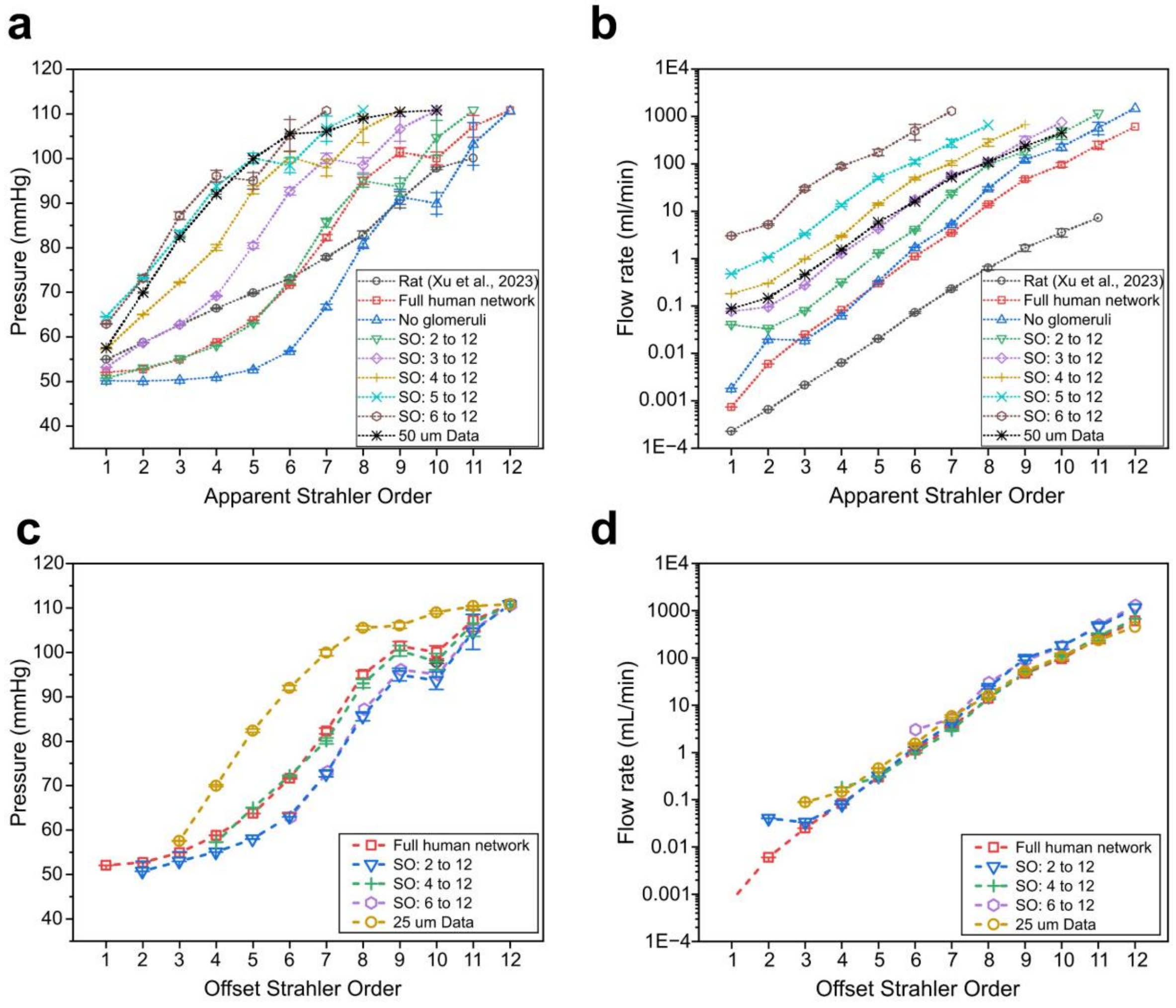


**SI Figure S2.** Simulation results showing the impact of vascular network completeness on renal hemodynamics. **(a)** Flow rate and **(b)** pressure distributions across apparent Strahler orders are shown for the full human network, networks with progressive pruning of low-order vessels (SO: 2 to 12 through SO: 6 to 12), and networks excluding glomeruli or based on 50 μm data alone. The plots illustrate how structural simplifications influence intrarenal flow and pressure patterns, particularly in lower Strahler orders. Rat simulation data from Xu et al. (2023)[9] are included for reference. **(c–d)** Pressure **(c)** and flow rate **(d)** plotted against offset Strahler order, enabling comparison of haemodynamics in equivalent anatomical vessels across networks with different levels of pruning.

In the main text (Figure 4c–d), we quantified the impact of incomplete vascular reconstruction by comparing pressure and flow values in vessels with the same Strahler order across networks with different levels of pruning (e.g. SO = 6). This comparison reflects how haemodynamic predictions change when only a limited number of vascular generations are resolved, as is typical in clinical imaging.

An alternative but complementary comparison can be made by examining the haemodynamics of *the same physical vessels* before and after pruning. When distal generations are removed, the Strahler order of the remaining vessels is redefined, such that a vessel with Strahler order $k$ in the pruned network corresponds to a vessel with Strahler order $k + n$ in the full network, where $n$ is the number of deleted generations. To enable direct comparison of equivalent vessels, we therefore applied a Strahler-order offset to the pruned networks, effectively reassigning their orders to match those in the full network.

Using this adjusted ordering, Supplementary Figure 2c–d shows pressure and flow rate as functions of the offset (equivalent) Strahler order. Under this comparison, differences between networks are reduced and exhibit a more complex, non-monotonic behaviour. For example, at an equivalent Strahler order of 6, mean pressure values are 72, 63, 73, and 62 mmHg for the full network and after pruning 1, 3, and 6 generations, respectively, with a corresponding value of 62 mmHg for the 50 μm network. Similarly, flow rates at the main renal artery are 607, 1021, 705, and 1052 mL/min, compared with 453mL/min in the 50 μm network.

This alternative analysis demonstrates that while incomplete reconstruction strongly alters haemodynamics when comparing vessels by nominal Strahler order, differences are smaller when comparisons are restricted to equivalent anatomical vessels. Together, the two analyses highlight distinct but complementary consequences of limited vascular resolution: changes in both the effective vascular hierarchy and the haemodynamic state of individual vessels.

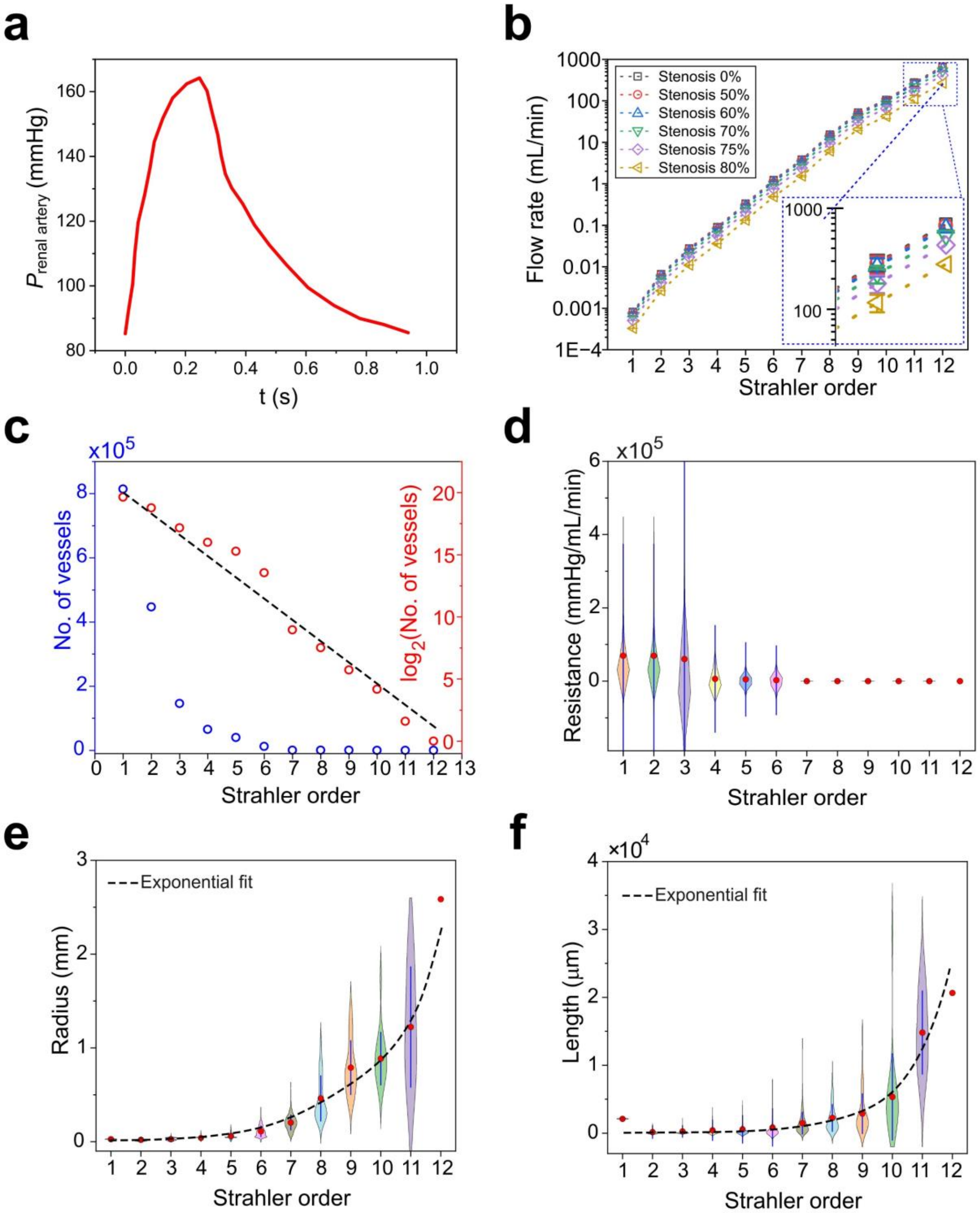


**SI Figure S3. Supplementary information on renal artery stenosis analysis. (a)** The pressure waveform applied at the renal artery inlet is based on *in vivo* measurements from a patient with renal artery stenosis, reported by Collard et al.. **(b)** Mean pressure and flow rate distribution across different Strahler orders under varying degrees of stenosis. **(c–f)** Physical and mechanical properties of the vasculature as a function of Strahler order, including vessel count **(c)**, vessel resistance **(d)** vessel radius **(e)**, and vessel length **(f)**.

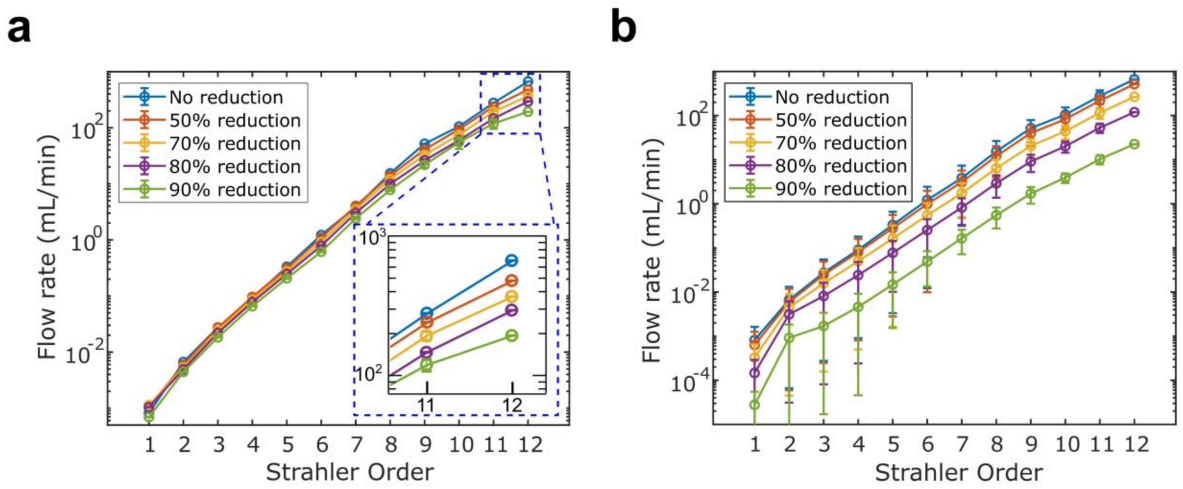


**Figure S4. Log-scale flow rates across Strahler orders under structural and functional rarefaction. (a)** Flow rate (log scale) as a function of Strahler order for structural rarefaction, corresponding to progressive removal of afferent arterioles and their glomeruli (0–90% reduction). **(b)** Flow rate (log scale) as a function of Strahler order for functional rarefaction, corresponding to progressive narrowing of afferent arteriole diameters (0–90% reduction).
